# Fine-grained modelling of ATP dependence of decision-making capacity in genetic regulatory networks

**DOI:** 10.1101/2023.11.16.567352

**Authors:** Rajneesh Kumar, Iain G. Johnston

## Abstract

Cellular decision-making is fundamental to life, from developmental biology to environmental responses and antimicrobial resistance. Many regulatory processes that drive cellular decisions rely on gene expression, which requires energy in the form of ATP. As even genetically identical cells can have dramatically different ATP levels, bioenergetic status can be an important source of variability in cellular decision-making. Existing studies have investigated this energy dependence but often use coarse-grained modelling approaches (which are not always readily connected to the underlying molecular processes of gene regulation). Here we use a fine-grained mathematical model of gene expression in a two-gene decision-making regulatory network to explore cellular decision-making capacity as energy availability varies. We simulate both a deterministic model, to explore the emergence of different cell fate attractors as ATP levels vary, and a stochastic case to explore how ATP influences the noisy dynamics of stochastic cell decision-making. Higher energy levels typically support increased decision-making capacity (higher numbers of, and more separated, cell states that can be selected), and the fine-grained modelling reveals some differences in behaviour from previous coarse-grained modelling approaches.

## Introduction

The supply of energy is fundamental to the out-of-equilibrium processes that maintain life, from physical motion and metabolism to gene expression and information processing (Mahmoudabadi et al. 2019; Flamholz, Phillips, and Milo 2014; Phillips and Milo 2009). ATP molecules serve as primary energy carriers in cells: many essential biological processes, including gene expression, metabolism, and transport, are regulated by ATP, which serves as a universal cofactor for cellular energy. The expression of genes, involving ATP-consuming transcription and translation, fundamentally determines a cell’s behaviour and its response to given conditions. The substantial ATP differences that exist across cells in a wide variety of organisms (Yaginuma et al. 2014; Takaine et al. 2019; Yoshida, Kakizuka, and Imamura 2016; De Col et al. 2017) can therefore give rise to substantial differences in gene expression and cell behaviour.

Gene regulatory networks (GRNs), where sets of genes influence each others’ expression, underlie a host of cell decision-making and information-processing mechanisms (Cortijo et al. 2019; Kalmar et al. 2009; Ozbudak et al. 2002; Blake et al. 2006; Fraser and Kaern 2009). A GRN may support a range of different expression profiles, each corresponding to a different cell state or cell fate. The response of GRNs to different environmental conditions, and the consequent changes in cell behaviour supported by GRN dynamics, give a cell “decision-making” capacity – that is, the ability to “decide” on a particular state of gene expression (Balázsi, Van Oudenaarden, and Collins 2011).

Changes in gene expression profiles, and thus in cell behaviour, may be driven by random chance as well as external stimuli. The discrete nature of molecular processes and the inherent randomness in reaction occurrences are commonly referred to as “intrinsic noise” at the molecular level of gene expression, where transcription and translation are carried out in environments with only a relatively small number of copies (Raj and Oudenaarden 2008; Shahrezaei and Swain 2008).. When molecular substances like DNA, mRNA, and regulatory proteins become scarce, there is intrinsic noise in biochemical processes (Elowitz et al. 2002; Ozbudak et al. 2002; Blake et al. 2006) that causes variability in their activity. Such variation may influence the dynamics of gene regulatory networks and cellular decision-making processes, causing unpredictability and uncertainty in the cellular response (Thattai and Van Oudenaarden 2001; Mitchison 2005; Walczak, Mugler, and Wiggins 2012; Thattai and Van Oudenaarden 2004; Acar, Mettetal, and Van Oudenaarden 2008). Intrinsic noise is now generally acknowledged to play a crucial role in gene regulatory networks (Kaern et al. 2005; Thattai and Van Oudenaarden 2001; Walczak, Mugler, and Wiggins 2012). It improves a single phenotype’s capacity to adapt to shifting environmental conditions (Thattai and Van Oudenaarden 2004; Acar, Mettetal, and Van Oudenaarden 2008) and can fundamentally shape cell decision-making (Guillemin and Stumpf 2020; Coomer, Ham, and Stumpf 2022).

A classic illustration of cellular decision-making in response to environmental stimuli is the Lac operon, where a bacterial cell carefully regulates lactose uptake and metabolism by regulating the operon’s expression in response to external factors (Vilar, Guet, and Leibler 2003). When lactose is in limited supply, the operon remains suppressed to conserve energy. However, lactose acts as an inducer when it is present, triggering the operon to be activated to promote lactose metabolism. With the support of this regulatory mechanism, the cell possesses the capability to efficiently adapt its processes of metabolism to changing environmental conditions.

Another example of bacterial decision-making reflects the influence of randomness. Persisters, stochastic variants in microbial populations, are characterized by their dormancy and high antibiotic tolerance. A drop in cytoplasmic ATP levels, along with stochastic changes in cell behaviour, is linked to the formation of persister cells across different species (Conlon et al. 2016; Shan et al. 2017; Zalis et al. 2019), contributing to their antibiotic tolerance and dormancy (although the universality of this link is debated (Braetz et al. 2017)).

The role of ATP in gene expression (Das Neves et al. 2010; Mahmoudabadi et al. 2019; Phillips and Milo 2009; Flamholz, Phillips, and Milo 2014; Kafri et al. 2016), and the fact that substantial cell-to-cell diversity can exist in ATP concentrations in different systems (Yaginuma et al. 2014; Takaine et al. 2019; Yoshida, Kakizuka, and Imamura 2016; De Col et al. 2017) means that ATP variability can lead to cell-to-cell variability in GRN dynamics and behaviour. This mirrors the ATP dependence of other biomolecular pathways (Gawthrop and Crampin 2017; 2014; Qian and Beard 2006) – for example, the detailed role of ATP in shaping the dynamics of signalling cascades has recently been illustrated using quantitative modelling grounded in systems biology and thermodynamics (Forrest et al. 2023). In GRNs, theoretical studies have shown that simplified GRN models (Karlebach and Shamir 2008), using coarse-grained descriptions of genes and gene expression processes, exhibit strong energy-dependent diversity in decision-making capacity (Johnston et al. 2012; Kerr, Jabbari, and Johnston 2019; Kerr et al. 2022). In such models of simple mutually repressing GRN motifs, increased ATP concentrations led to more attractor states (that is, distinct stable states of gene expression to which other states evolve over time), and hence to more decision-making capacity in cells. Lower ATP concentrations reduced the attractors supported to the point where only a single mode of behaviour was permitted, compromising decision-making ability. Assuming that ATP concentration scales the rate of gene expression processes (Das Neves et al. 2010), this behaviour is compatible with classic studies on the attractor structure of model GRNs (Gardner, Cantor, and Collins 2000; Huang et al. 2007), which typically find that increased rates of gene expression support more, and more stable, dynamic attractors.

Although informative, many of these studies used coarse-grained models of gene expression (Karlebach and Shamir 2008; Caranica and Lu 2023), neglecting differences between RNA and protein (using a single, combined “gene” variable to track expression) and a heuristic Hill function model for gene interactions. Further, the dynamics of gene expression are fundamentally stochastic, but the investigation of ATP influence on stochastic models of GRN dynamics are limited. The question therefore remains open: do more realistic models of GRN behaviour also show energy dependence of decision-making?

To address this, we implemented a GRN model that explicitly tracks DNA, RNA, and protein concentrations, where dimerised proteins act to control gene expression in a mutually antagonistic motif, from (Kim and Gelenbe 2012) and ultimately deriving from (Gardner, Cantor, and Collins 2000), with parameters and reactions all corresponding to empirically measurable observables. We explored conditions under which this model demonstrates decision-making capacity (the appearance of more than one attractor) and how this capacity depends on ATP.

## Methods

### Model description

We use the switch model from Kim & Gelenbe (Kim and Gelenbe 2012), following the classical toggle switch model in (Gardner, Cantor, and Collins 2000), but remove the delay aspect of translation to simplify the system. The model includes two genes i and j, the protein products of which dimerise. The dimer of one gene inactivates the promoter of the other (Fig. 1). In other model variants, we introduce further (or fewer) polymerization processes, so that the polymer of size n is the regulatory factor influencing gene expression, where n range from 1 (monomer regulation) to 4 (tetramer regulation). We will mainly focus on the well-studied n=2 case.

**Figure 1.**
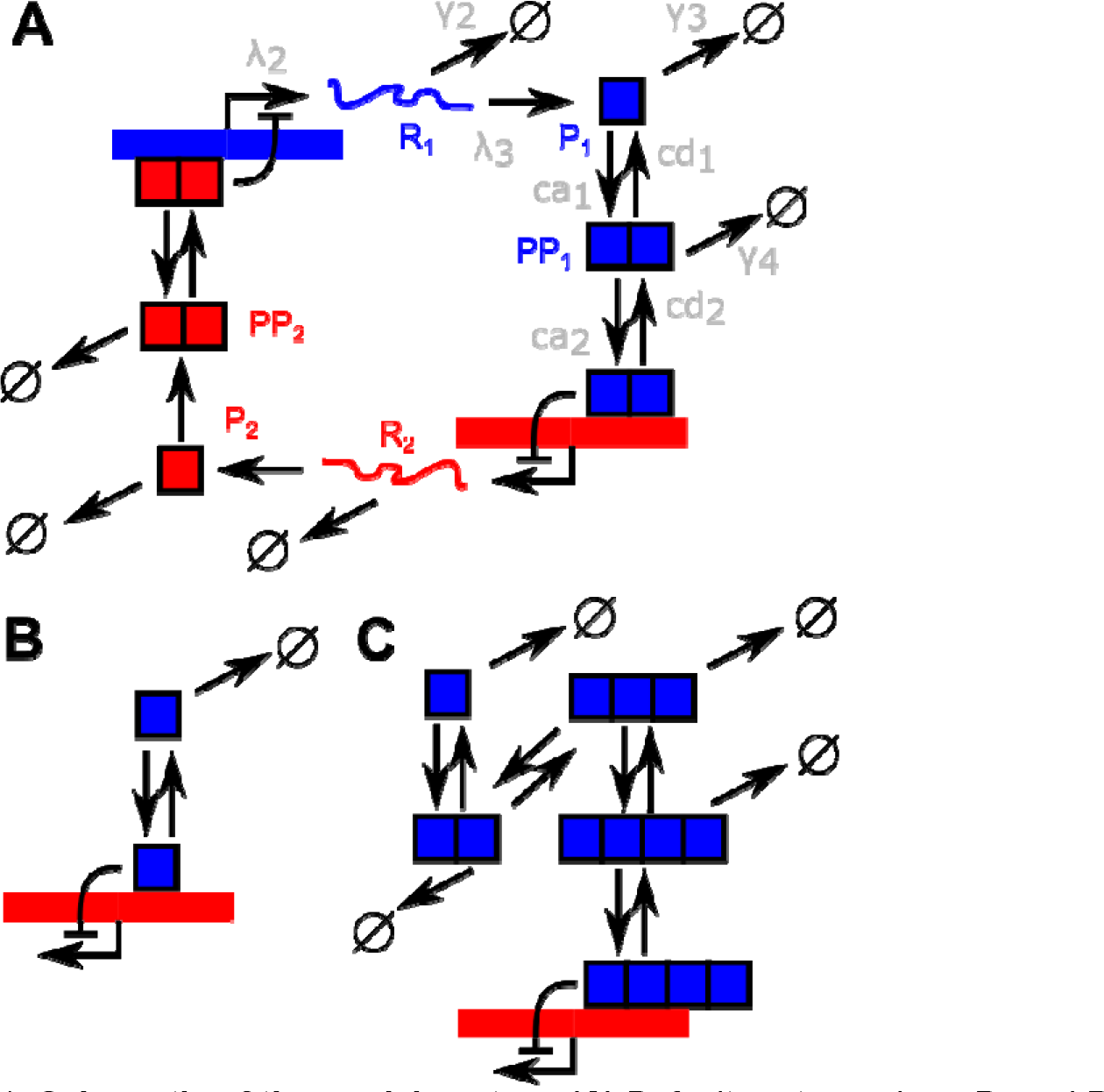
Schematic of the model system. **(A)** Default system, where P_1_ and P_2_ are expressed and their dimers repress each others’ expression. For clarity, rates are only shown on processes involving the first product; they apply symmetrically to the other. λ_2_ and λ_2_ will be scaled by model ATP concentration to capture ATP dependence of transcription and translation. **(B)** Subset of simplified system without polymerization, where the monomer acts as the regulatory factor. **(C)** Subset of more complicated system where oligomerization proceeds up to the homotetramer, which acts as the regulatory factor.

We use Kim & Gelenbe’s (Kim and Gelenbe 2012) parameterisation, itself taken from the biological literature (Paulsson 2005; Thattai and Van Oudenaarden 2001; Bratsun et al. 2005; Goeddel, Yansura, and Caruthers 1977; Buchler, Gerland, and Hwa 2005). We set transcription factor concentrations equal to a unit constant, reflecting a constant constitutive level of expression, and we initialize the system with Pro_i_ = 1 (one promoter for each gene), initial protein levels P_1_ = P_1_*, P_2_ = P_2_*, and all other species at zero concentration. We will model ATP dependence by scaling gene expression processes by a constant given by a model ATP value (Kerr, Jabbari, and Johnston 2019; Kerr et al. 2022), capturing the dependence of transcription and translation rates on ATP concentration (Das Neves et al. 2010).

### Differential equation model

We explored both deterministic and stochastic instances of this system. The state variables in our system are illustrated in Fig. 1: p_i_ is the monomer protein of gene i and pp_i_ is the dimer of this protein, pro_i_ is the promoter for gene i, propp_i_ is promoter j bound by dimer pp_i_, and rna_i_ is mRNA for gene i. For the deterministic picture we used the governing ordinary differential equations to capture a mass action model of the processes in Table 1:

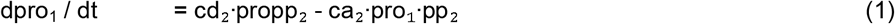

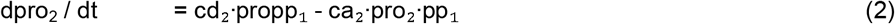

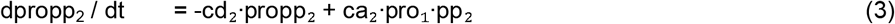

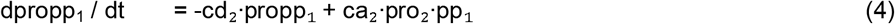

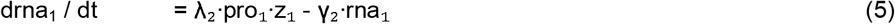

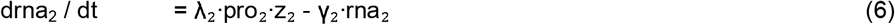

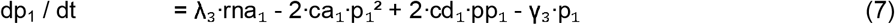

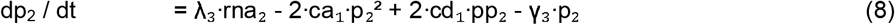

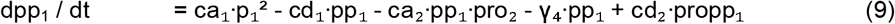

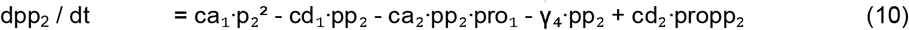

with the following initial conditions

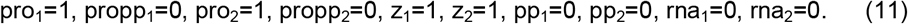

**Table 1.** Processes in the model.

| Process | Reaction | Rates (default parameterization value in s<sup>-1</sup>) |
| --- | --- | --- |
| Transcription from active promoter | $\text{Pro}_i + \text{TF}_i \rightarrow \text{Pro}_i + \text{TF}_i + \text{R}_i$ | $\lambda_2$ (0.0067) |
| Translation | $\text{R}_i \rightarrow \text{R}_i + \text{P}_i$ | $\lambda_3$ (0.0474) |
| Dimerisation/Dedimerisation | $\text{P}_i + \text{P}_i \leftrightarrow \text{PP}_i$ | $\text{ca}_1 / \text{cd}_1$ (0.0023 / 0.00023) |
| Dimer binding/unbinding to promoter * | $\text{Pro}_i + \text{PP}_j \leftrightarrow \text{Pro}_i \text{PP}_j$ | $\text{ca}_2 / \text{cd}_2$ (0.1038 / 1.04) |
| RNA degradation | $\text{R}_i \rightarrow 0$ | $\gamma_2$ (0.00023) |
| Protein degradation | $\text{P}_i \rightarrow 0$ | $\gamma_3$ (0.00077) |
| Dimer degradation | $\text{PP}_i \rightarrow 0$ | $\gamma_4$ (0.00058) |
| **Model variant processes** (only one promoter interaction * is present in any model instance) |  |  |
| Monomer binding/unbinding to promoter * | $\text{Pro}_i + \text{P}_j \leftrightarrow \text{Pro}_i \text{P}_j$ | $\text{ca}_2 / \text{cd}_2$ (0.1038 / 1.04) |
| Trimerisation/detrimerisation | $\text{P}_i + \text{PP}_i \leftrightarrow \text{PPP}_i$ | $\text{ca}_1 / \text{cd}_1$ (0.0023 / 0.00023) |
| Trimer binding/unbinding to promoter * | $\text{Pro}_i + \text{PPP}_j \leftrightarrow \text{Pro}_i \text{PPP}_j$ | $\text{ca}_2 / \text{cd}_2$ (0.1038 / 1.04) |
| Trimer degradation | $\text{PPP}_i \rightarrow 0$ | $\gamma_4$ (0.00058) |
| Tetramerisation/detetramerisation | $\text{P}_i + \text{PPP}_i \leftrightarrow \text{PPPP}_i$ | $\text{ca}_1 / \text{cd}_1$ (0.0023 / 0.00023) |
| Tetramer binding/unbinding to promoter * | $\text{Pro}_i + \text{PPPP}_j \leftrightarrow \text{Pro}_i \text{PPPP}_j$ | $\text{ca}_2 / \text{cd}_2$ (0.1038 / 1.04) |
| Tetramer degradation | $\text{PPPP}_i \rightarrow 0$ | $\gamma_4$ (0.00058) |

### Oligomerisation model

Let *x* be a monomer that may be capable of self-oligomerisation. Let q_*0*_ be an inactive promoter and q_*1*_ be an active promoter. Let *r* be a molecule, which will be an oligomeric state of *x*, which activates the promoter. We have the following system

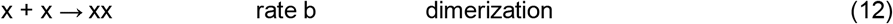

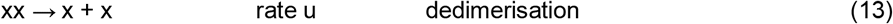

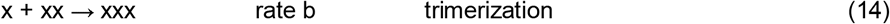

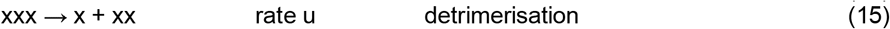

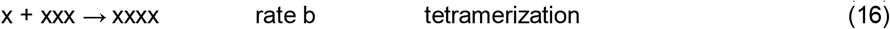

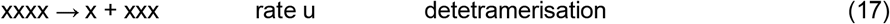

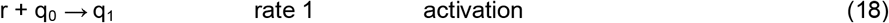

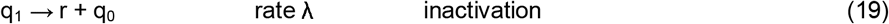

We consider three instances:

i. Active monomer: none of processes 12-17 occur, *r* is *x*
ii. Active dimer: processes 12-13 occur, *r* is *xx*
iii. Active tetramer: processes 12-17 occur, *r* is *xxxx*

We start the system with *x* = *x*_0_, *q*_0_ = 1, all other concentrations zero. We ask what happens to steady-state *q*_1_ (proportion of activated promoter) as *x*_0_ varies. For the illustrative example in the text we use *b* = 0.05, *u* = 1 (so oligomerisation is relatively infrequent), *λ* = 0.001 (so inactivation is infrequent).

### Numerical implementation

The system exhibited dynamics which made numerical solution somewhat awkward with some solvers. Specifically, the system quickly relaxes to a set of states where future time behaviour is very slow (Supp. Fig. 1). Several solvers misassessed this slow behaviour as stationarity and did not proceed to the true steady-state solution. To avoid such artefacts, we used a hard-coded forward Euler solver with a convergence criterion defined over a time window, and confirmed that an LSODA solver in R (Soetaert, Petzoldt, and Setzer 2010) also successfully negotiated this slow-changing dynamic. For the stochastic simulation, Gillespie’s stochastic simulation algorithm (Gillespie 1977) was used to explicitly simulate trajectories from the processes in Table 1.

The numerical simulations were performed using custom code in C for speed, with some checking using *deSolve* (Soetaert, Petzoldt, and Setzer 2010) in R. For data handling and visualization we used R libraries *ggplot2* (Wickham 2011), *reshape* (Wickham 2007), and *ggpubr* (Kassambara 2020). All code is freely available at https://github.com/StochasticBiology/energy-decisions.

## Results

### Parameter values supporting decision-making capacity in the deterministic model

Within the context of this model, cellular decision-making capacity is defined as the support of distinct cell states defined by protein concentrations (resembling different *in vivo* cellular phenotypes). If the system always converges to a single state, there is no capacity to “make a decision” and adopt a different phenotype according to context. By contrast, if several distinct cellular states are supported by the system, the ability to “decide” between them is supported. States to which a system converges over time are called “attractors”, and the set of initial states from which an attractor is eventually reached is called an “attractor basin”.

To identify parameterisations of the model that led to non-trivial decision-making capacity, we conducted a stepwise sweep through parameter values over orders of magnitude and looked for those parameterizations that resulted in an attractor basin structure with more than one attractor. Our parameter sweep centred on the biologically-derived parameters from (Kim and Gelenbe 2012) and varied each over surrounding orders of magnitude. The support for more than one attractor demonstrates that the system can exhibit different steady states (behaviours) in response to different stimuli or environmental conditions.

During this investigation we found that the dynamics of the system exhibited rather stiff characteristics: the system rapidly evolves to occupy a region of state space then slowly evolves further within this set of states before finally reaching an attractor. In several instances the system showed non-monotonic behaviour with time in all state variables (Supp. Fig. 1), with coincident turning points resembling a “false convergence” to a steady state. We tested for convergence over a longer time period to avoid numerical issues with the ODE solvers involved (Methods).

The results of this parameter sweep for the default system where dimers are the active regulatory molecules (Fig. 1A) are shown in Fig. 2. We found in particular that increased protein degradation (γ_3_) and decreased TF-DNA dissociation (cd_2_) supported a tripartite system where one central attractor (with equal protein levels) was flanked by two attractors (high pro_1_ and high pro_2_ respectively). In cases where this tripartite structure was not present, the system was typically characterised by relaxation towards a single attractor. Other combinations of parameters supporting non-trivial attractor structure likely exist, but we focussed on these two cases first as the most directly connected to biological values.

**Figure 2.**
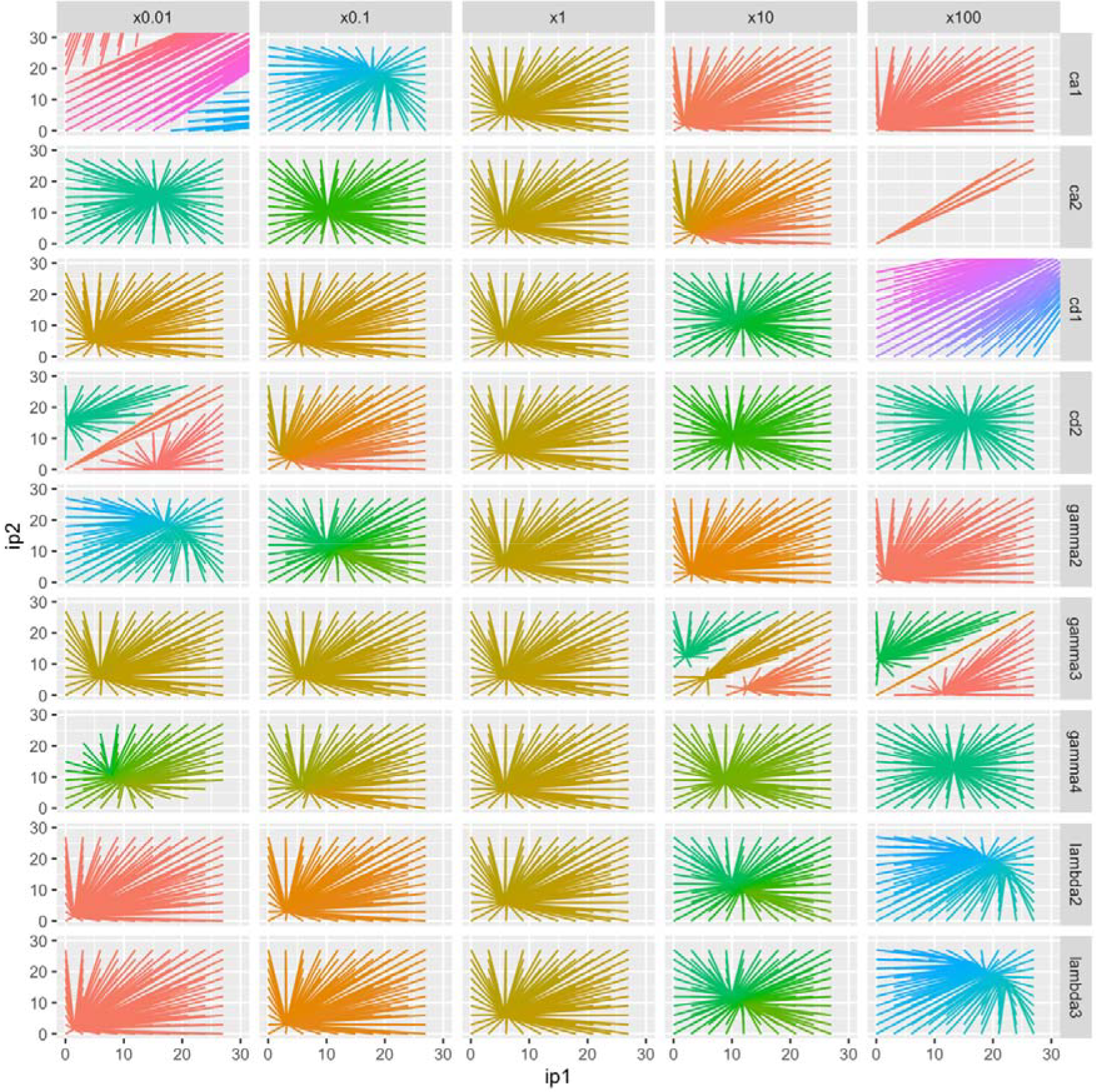
Attractor basin structure with orders-of-magnitude scans through parameters. Each line describes how the given initial condition (ip_1_, ip_2_) evolves to a long-term position in the (p_1_, p2) space of protein expression. Rows correspond to different model parameters being varied; columns correspond to the scaling applied to each parameter. Colours are assigned based on the coordinates of the long-term state. Tripartite attractor structure is caused by lower values of TF unbinding rate cd_2_ and higher values of protein degradation rate γ_3_ (and, on a different scale, lower values of dimerization rate ca_1_). Missing lines represent situations in which the system failed to converge over the simulated timescale.

### Effect of required oligomerization state of regulatory factor

The default structure of the model has the dimer of the protein product as the active regulatory factor. Following previous work exploring the effect of different degrees of model cooperativity on decision-making capacity (Kerr, Jabbari, and Johnston 2019), we next asked how the required oligomerization state of the regulatory protein product influenced decision-making capacity. Supp. Fig. 2 shows the more general results where gene expression is regulated by monomer, trimer, and tetramer protein products. When the monomer is the active factor, most parameterizations lead to more trivial attractor basin structure. This mirrors previous work (Kerr, Jabbari, and Johnston 2019) where lower Hill coefficients in a coarse-grained model led to decreased decision-making capacity – a result of the more linear, less switch-like regulatory interactions supported in the absence of cooperativity.

More unexpectedly, increasing the required oligomerization state of the regulatory factor to trimeric or tetrameric respectively showed little change and a negative change in decision-making capacity (Supp. Fig. 2). The required oligomerization state of the transcription factor has the same quantitative influence on the microscopic system as an increased Hill coefficient has on the coarse-grained system – broadly increasing the non-linearity of the increasingly step-like response (Supp. Fig. 3; see (Hernández-García and Velázquez-Castro 2023)). In coarse-grained modelling, increasing Hill coefficients supported more attractor diversity (Kerr, Jabbari, and Johnston 2019). In this microscopic case, there is a limit of oligomerization beyond which the increase in decision-making capacity stops and then reverses.

### Influence of protein degradation and dimer-DNA dissociation on decision-making ability in the deterministic model

Following this exploration, we first investigated the effect of increasing protein degradation rate γ_3_ in the model. The attractor basin structure in this case shows re-entrant behaviour, with a single attractor appearing for low and high degradation values, and tripartite structure appearing for intermediate values (Fig. 3A).

**Figure 3.**
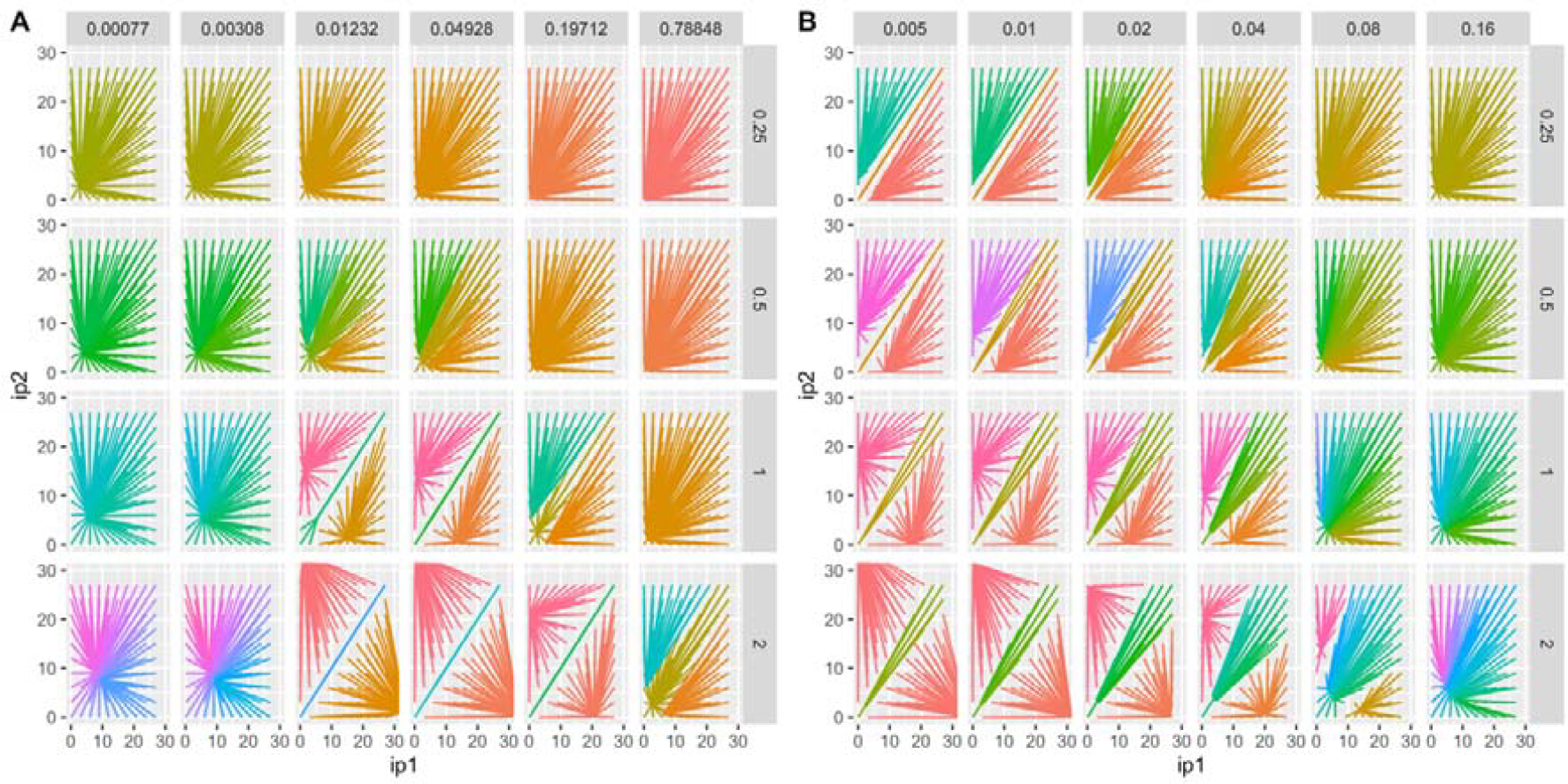
Attractor basin structure with degradation and dissociation rates, and model ATP variability. Attractor structure with **(A)** varying protein degradation rate γ_3_ (columns) and ATP (rows); **(B)** varying TF unbinding rate cd_2_ (columns) and ATP (rows). As in Fig. 1, each line describes how the given initial condition (ip_1_, ip_2_) evolves to a long-term position in the (p_1_, p2) space of protein expression. Colours are assigned based on the coordinates of the long-term state.

We next decrease the DNA-protein dissociation rate cd_2_, so that the dimeric transcription factors stay bound to their opposing promoter regions for longer. This increased “memory” induces separation of the attractors of the system. Decreasing cd_2_ reduces the size of the central attractor, so that the stability of the non-zero attractor relative to the central attractor increases as the value of cd_2_ rises.

### ATP modulation of transcription and translation controls decision-making ability in the deterministic model

Having characterized the behaviour of decision-making capacity with some key parameters of the model, we next addressed our central question – how energy levels influence decision-making capacity. To this end, we modulate transcription and translation rates λ_2_ and λ_3_ in the model by a factor proportional to ATP concentration, and allow this concentration to vary. The true relationship between ATP concentration and these rates has been modelled as sigmoidal (Das Neves et al. 2010; Kerr, Jabbari, and Johnston 2019) but we effectively employ a linear approximation to the centre of this sigmoidal behaviour for simplicity here.

We found that increasing model ATP concentration generally enhanced decision-making capacity in model cells, reflected in the appearance of more attractor basins and increased separation between them (Fig. 3). Higher ATP concentrations – and hence higher rates of gene expression – typically led to increased separation of the cell states where one protein was present at a high level and the other at a low level. In tandem, the width of the central attractor basin corresponding to balance protein levels also increased. This behaviour mirrors that observed in coarse-grained empirical models (Johnston et al. 2012; Kerr, Jabbari, and Johnston 2019), confirming that this behaviour is not an artefact of the Hill function modelling approach and is expected to arise from the dynamics of individual regulatory interactions.

### Stochastic modelling of cell behaviour

The above deterministic modelling suggested that increasing cellular ATP levels supports more distinct cell states, corresponding to increased decision-making capacity. We next asked whether this behaviour was supported by a yet more detailed modelling approach (Karlebach and Shamir 2008) – a discrete stochastic model considering the copy numbers and interactions of each individual molecule in the system. To this end, we used Gillespie’s stochastic simulation algorithm (Gillespie 1977) to simulate trajectories of the system under different parameterisations corresponding to different cellular ATP levels.

Typical stochastic trajectories of the system are shown in Fig. 4A-B. Here, because random fluctuations in copy number can occur, the system does not remain forever in one attractor. Instead, fluctuations can drive the system out of the basin of one attractor and into another – corresponding to a “switch” from, for example, a high p_1_, low p_2_ state into a low p_1_, high p_2_ state. In cases where distinct cell states do not exist, the behaviour of the system simply corresponds to random fluctuations of proteins around the sole attractor. In cases where distinct states do exist, switching between states is observed – with a frequency that decreases as separation of the distinct states increases. Fig. 4C shows this behaviour more explicitly – that is, for higher ATP levels, the distribution of switching times between distinct basins has more density at longer times (with some remaining density at short times due to transient behaviour around switches), corresponding to more distinct and stable cellular phenotypes, and hence more robust decision-making capacity. Taken together, the stochastic and deterministic models of this system therefore agree (internally and with existing work) that increasing ATP supply supports more, and more stable, capacity for cellular decision-making through GRN dynamics.

**Figure 4.**
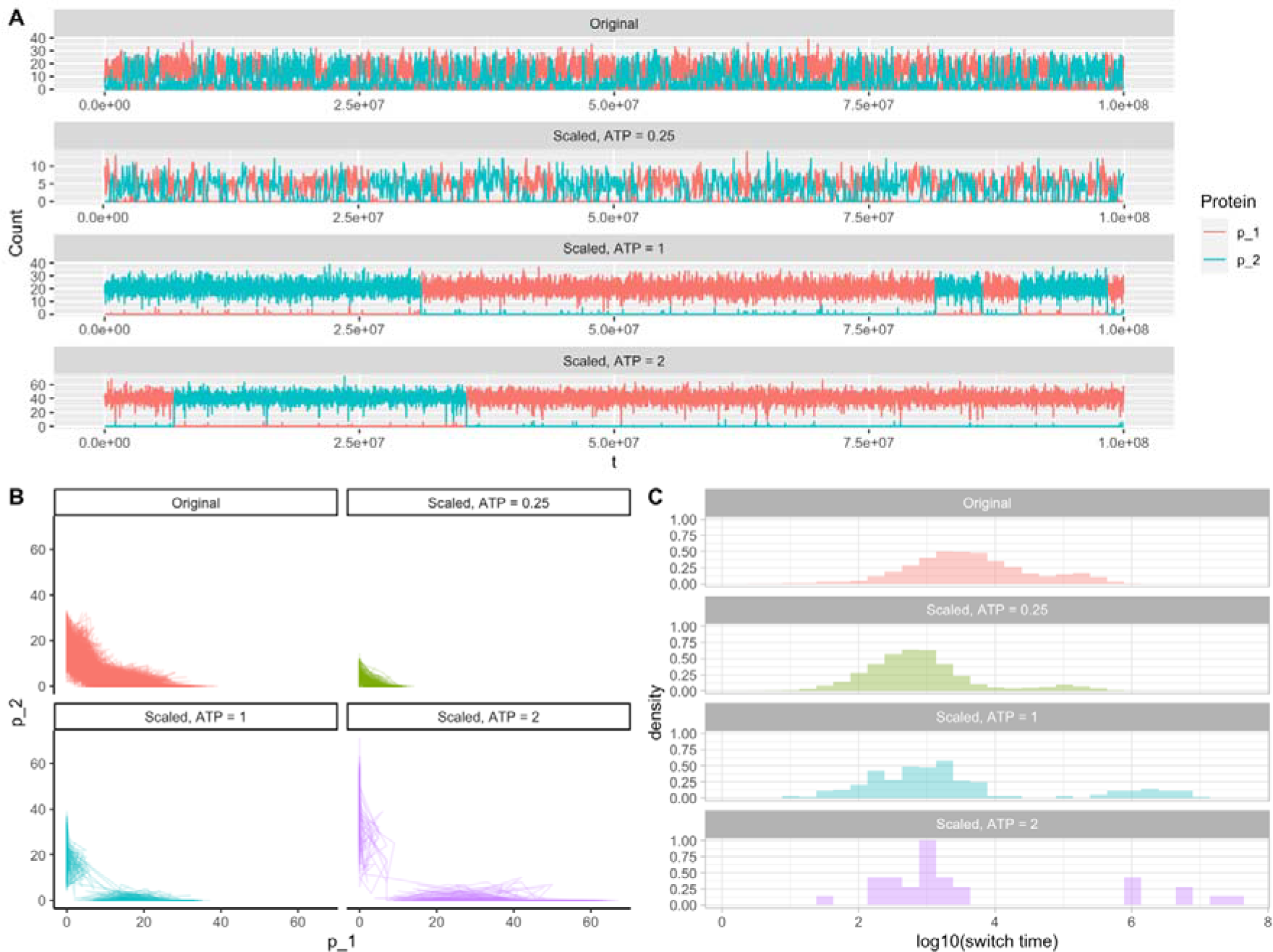
Stochastic dynamics of the model system. **(A)** Representative trajectories of protein expression levels (p_1_ and p_2_) under different ATP conditions. **(B)** Trajectories of the system in (p_1_, p_2_) space over time. **(C)** Distribution of (log) switching times (the time taken for the dominant protein in the system to change). Experimental labels: Original – original parameterization from (Kim and Gelenbe 2012), with TF unbinding rate cd_2_ = 1.04. Scaled – adapted parameterization with cd_2_ = 0.04 and model ATP levels as given.

## Discussion

We have shown that fine-grained mechanistic models, both ODE-based and stochastic, for GRN-based cell decision-making exhibit strong dependence on ATP supply, supporting results from more coarse-grained modelling (Kerr, Jabbari, and Johnston 2019; Kerr et al. 2022; Johnston et al. 2012) and from different specific systems (Forrest et al. 2023). According to (Kerr, Jabbari et al. 2019), increasing intracellular energy levels can increase the number of supported stable phenotypes, hence enhancing the capacity for decision-making. We find qualitatively similar behaviour here, with quantitative details dependent on the polymerization structure of the transcription factor governing regulation within the GRN.

Classical studies on GRNs supporting various switch-like behaviours (Gardner, Cantor, and Collins 2000; Huang et al. 2007) consistently observe that increasing rates of gene expression stabilize and separate the attractor basins of a model. This picture aligns with our findings and those in (Kerr, Jabbari, and Johnston 2019; Johnston et al. 2012), where the effect of increasing ATP supply is captured by increasing such rates of expression. The classic (Gardner, Cantor, and Collins 2000) also observes dependence on the steepness of the regulatory interaction between genetic components – specifically, that higher Hill coefficients (steeper switching) expand and stabilize an intermediate attractor basin where both genes are expressed. This expanded attractor structure agrees with theory using coarse-grained Hill functions (Kerr, Jabbari, and Johnston 2019). However, the fine-grained modelling we use here suggests some nuance to this picture – specifically that steeper switching (captured here by increasing the required polymerization of the regulatory factor) only stabilizes expanded attractors up to a point, and that further increases limit decision-making capacity.

Our modelling approach differs from the original instance in (Kim and Gelenbe 2012) (following (Gardner, Cantor, and Collins 2000)) by the removal of the delay term in the translation process. We made this change for simplicity of interpretation and to facilitate comparison with existing modelling approaches. One consequence of this change is that the parameterization from the original paper does not demonstrate dramatic bistability in our model; changes to parameters (most directly, either through protein degradation or through TF unbinding) are needed to separate attractor basins in the system’s behavioural profile. Exploration of the effects of jointly varying parameters in this model, perhaps through sensitivity analysis regarding the number of distinct attractors as a summary of the system’s response, would be interesting to further characterize the detailed behaviour of this model. In parallel, exploration of different initial conditions reflecting diversity in the non-protein elements (particularly mRNA) may be of interest to explore differences in transient behaviour of the system.

The fundamental importance of ATP of course means that it influences other cellular processes than gene expression. Cells and organisms respond to changes in ATP supply in a wide variety of ways. In humans, for example, ATP depletion can adversely affect essential cellular processes, potentially leading to organ failure and cellular dysfunction (Johnson, Jinnah, and Kamatani 2019). Numerous disorders have been scientifically linked to the occurrence of cellular ATP shortages. Some of these disorders include mitochondrial diseases (Johnson, Jinnah, and Kamatani 2019), metabolic disorders, neurodegenerative diseases (Cisneros-Mejorado et al. 2015), and myopathies. In bacteria, a decline in intracellular ATP production contributes to the formation of persister cells in different species (Conlon et al. 2016; Shan et al. 2017; Zalis et al. 2019) – itself a decision governing by changing gene expression profiles. The general finding that limited ATP supply compromises cellular decision-making through its influence on gene expression must be interpreted in parallel with these facts that ATP responses themselves can induce cellular decisions, and the link between these ATP response decisions and the cell’s energetic ability to make them will be a target for further research.

## Acknowledgments

This work was supported by the Trond Mohn Foundation [project HyperEvol under grant agreement No. TMS2021TMT09], through the Centre for Antimicrobial Resistance in Western Norway (CAMRIA) [TMS2020TMT11]. This project has received funding from the European Research Council (ERC) under the European Union’s Horizon 2020 research and innovation programme [grant agreement No. 805046 (EvoConBiO)].

## Supplementary Information

**Supplementary Figure 1.**
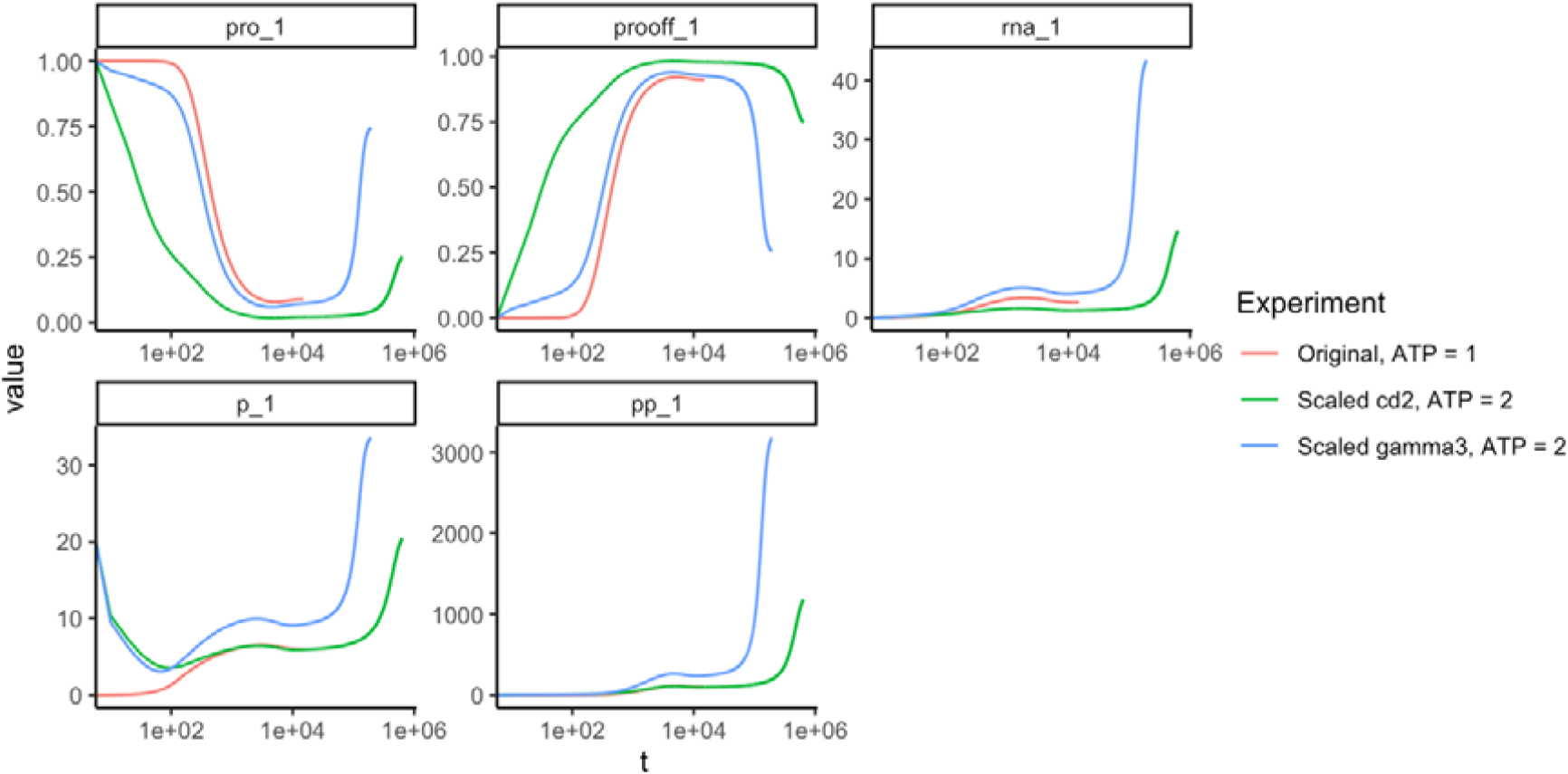
Example time series behaviour of the system. State variables in the system under three different profiles: default parameterization and symmetric initial conditions; model ATP = 2 and lower TF unbinding rate cd_2_ with asymmetric initial conditions; model ATP = 2 and lower protein degradation rate γ_3_, and asymmetric initial conditions. In all cases the system undergoes a long period of very slow change, often passing through turning points before finally converging (end of each trajectory).

**Supplementary Figure 2.**
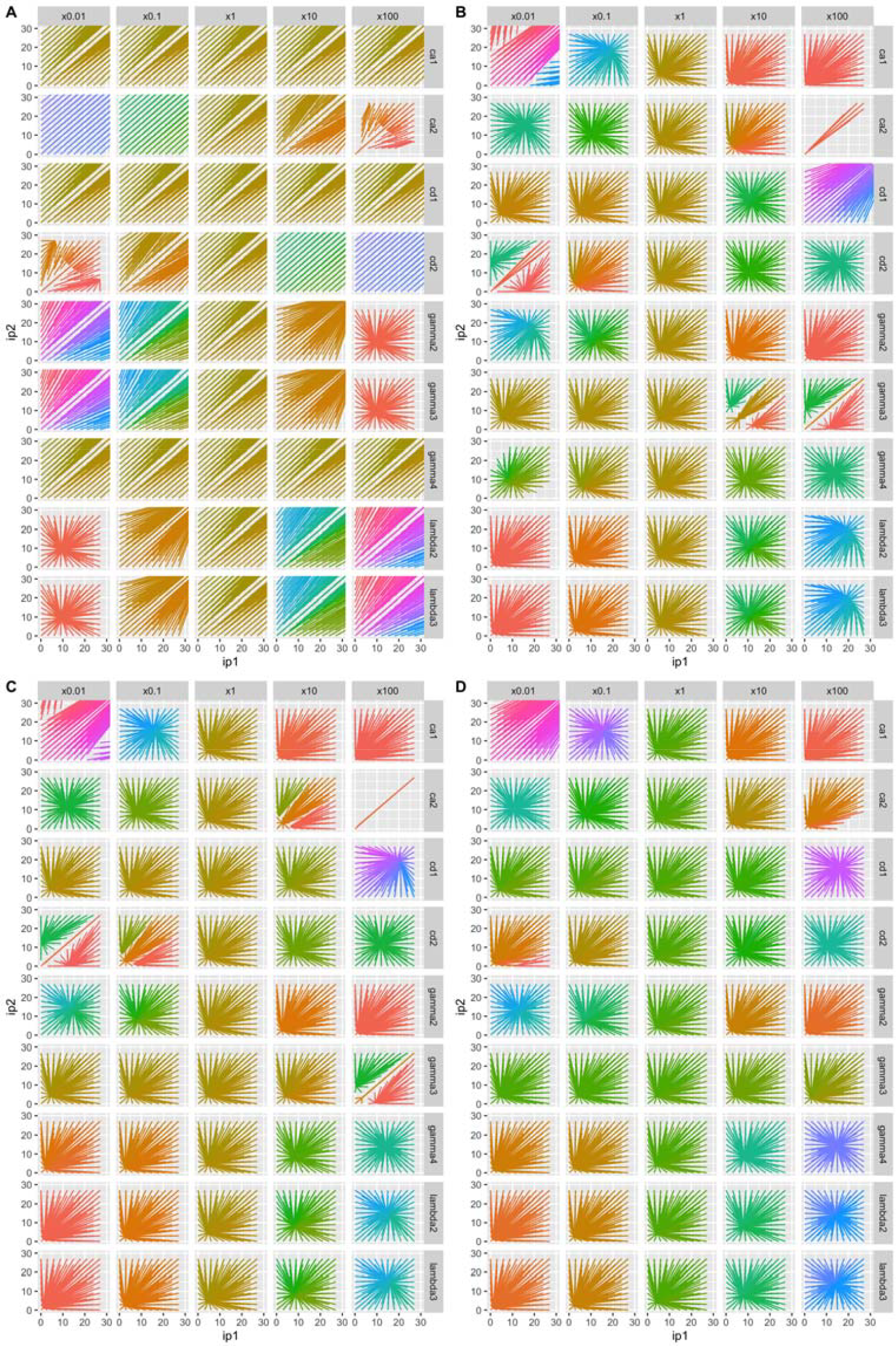
Attractor basin structure upon varying model parameters for different regulatory structures. Plots, as in Fig. 2, show evolution of the protein level in the system from different initial conditions, as parameters (rows) are varied over scales (columns). Colours are assigned based on the coordinates of the long-term state. The four panels **(A, B, C, D)** correspond to n=1, 2, 3, 4; hence regulation imposed by monomer, dimer, trimer, tetramer protein products.

**Supplementary Figure 3.**
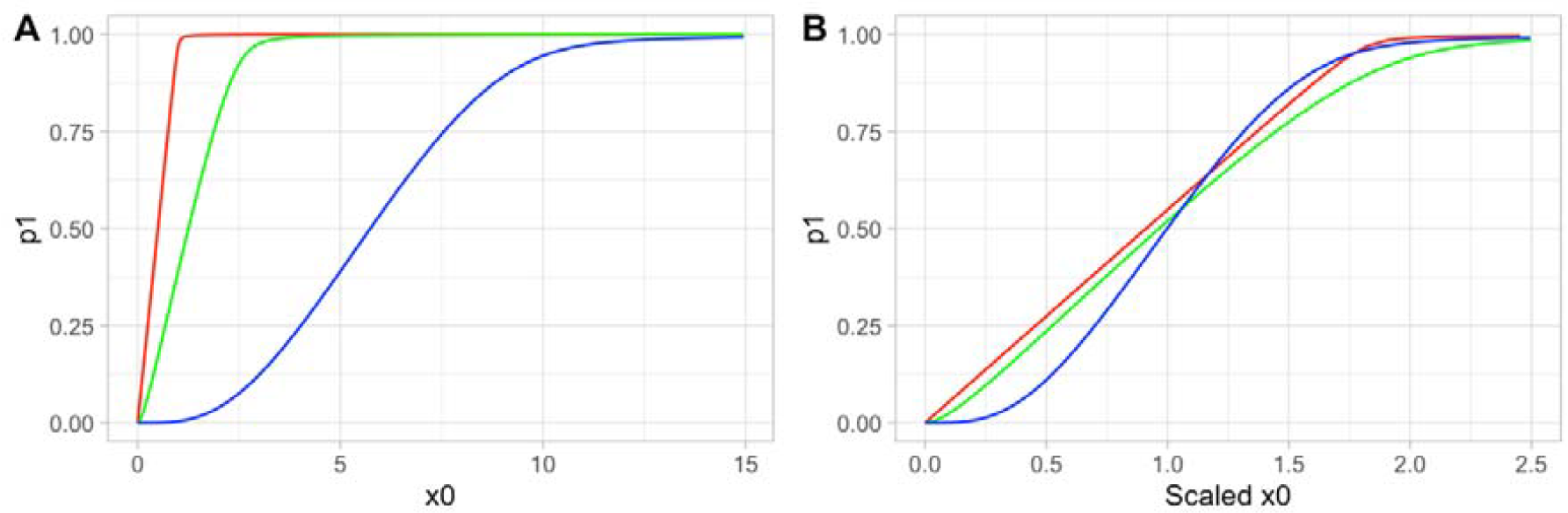
Promoter response for different transcription factor oligomerisations. **(A)** Profile of activated promoter *p1* with initial monomer concentration *x0*. **(B)** Same profile but with *x0* scaled in each case so that the 50% activation points for each curve overlap. Red, active monomer; green, active dimer; blue, active tetramer.

